# Consolidation of a recurrent choice pattern into a fixed action schema requires the orbitofrontal cortex

**DOI:** 10.1101/2025.05.08.652969

**Authors:** Ryo Tonami, Sho Aoki, Kentaro Abe

## Abstract

Our actions are crystallized by repetition, warranting the efficiency, reproducibility and stability of decisions. Unlike a habit that is formally defined as insensitive to reward value, such persistency in choice can still be shaped and executed in a goal-directed manner, which underlies many volitional behaviors in animals. Yet, largely due to the absence of an established procedure that accesses *non-habitual* but persistent patterns of choice in animal models, empirical findings that identify its neural substrates are lacking. Using a cost-benefit conflicting decision paradigm in rodents, we here introduce an experimental procedure involving extensive decision opportunities in a fixed contingency, whereby mice acquire a recurrent pattern of choice and shape it into a fixed schema of goal-directed action to guide behavior. A subsequent functional screening of neural circuitry by chemogenetic inhibition reveals the orbitofrontal cortex (OFC) as a candidate for this form of choice. Notably, we found that OFC plays a specific role in embedding the recurrent choice pattern into the fixed schema for action selection, but not in merely defining conflicting choice. Consistently, we uncover that OFC ablation leaves animals’ ability intact to make a conflicting choice, but selectively disrupts the formation of the fixed action schema. These findings suggest that OFC is crucial to ensure persistency in action selection by solidifying extensive history of recurrent choices into a firmly established schema of action selection, providing novel insights into how the stability and persistency of behavior are held in the brain.

## Introduction

Animals learn volitional actions by experience and their repetitions crystallize acquired behaviors ^1,2^. Such refinement in actions and decisions often leads to automaticity of behaviors, ensuring their efficiency, reproducibility and stability ^3–8^. One way to enable behavioral automaticity is the formation of habits that are stable but inflexible ^9–13^, contrasting to more flexible goal-directed behaviors ^14–17^. More formally, whether an agent’s behavior is habitual or goal-directed is defined by its sensitivity to dissociation of action-outcome contingency under reward devaluation or contingency degradation ^12,18–20^. This distinction has been commonly accepted and has provided a framework of how our behavior becomes stable, automated and efficient.

While this has been influential to behavioral control theories, it may overlook a more nuanced form of action selection animals do, especially those that inherit both features of habitual and goal-directed behaviors. Several studies have indeed suggested that animals demonstrate stereotypical behavioral patterns or sequential actions via their extensive repetition, but these are still goal-oriented ^11,21–23^. Similarly, psychological studies have agreed that human behaviors entail perseverance and persistency to an acquired pattern of choice, characterized by tendency to repeat a similar choice based on prior experiences ^24–27^. Therefore, it is no doubt that animals including humans acquire and execute stereotypical actions in goal-directed choice behavior. One way to achieve this is to consolidate recurrent choices we make repeatedly into one’s fixed action schema for guiding goal-directed behavior ^28,29^, such that action selection becomes more efficient, stable, and resistant to change. Notably, prior studies have suggested the involvement of frontal cortices ^30–32^, basal ganglia ^33–35^ and/or hippocampus ^36,37^ in representing stabilized action selection or refined sequences, showing candidate brain regions where fixated action schemas can be stored. Because of the limitation of empirical findings, however, neural substrates that play a causal role in shaping them remain largely unknown.

Inspired by and adopted from the cost-benefit conflicting decision task in rodents ^38,39^, we here introduce a novel experimental procedure by which mice execute a recurrent pattern of goal-directed choice and consolidate it into their fixed schema of action selection. Combining this paradigm with chemogenetics-based functional screening and neuronal ablation, we identified that the orbitofrontal cortex (OFC) plays a critical role in shaping the recurrent choice pattern into the established action schema to guide behavior, ensuring persistency in action selection. Our findings provide crucial insights into how the brain shapes the fixed pattern of action and how this function can be compromised in diseases.

## Results

### Mice acquire a recurrent choice pattern and shape it into a fixed action schema

Based on a well-established cost-benefit decision task in rodents ^38,39^, we designed an experimental procedure that allowed the animals to form a recurrent pattern of choice and consolidate it as their fixed schema for action selection (Fig. 1A-C). This procedure requires four phases in consecutive behavioral sessions from continuous reinforcement schedule (CRF, Fixed-ratio 1), a Reward-Value based choice session (Value session), a Reward-Value/Effort conflict session (Conflict session), and one-day Value session after extensive training on the Conflict sessions (Post-Conflict Value session) (Fig. 1C). Each session was composed of three blocks, where a block began with a forced choice trial on each of small and large reward choices, followed by 10 trials of free choice (Fig. 1D).

**Figure 1.**
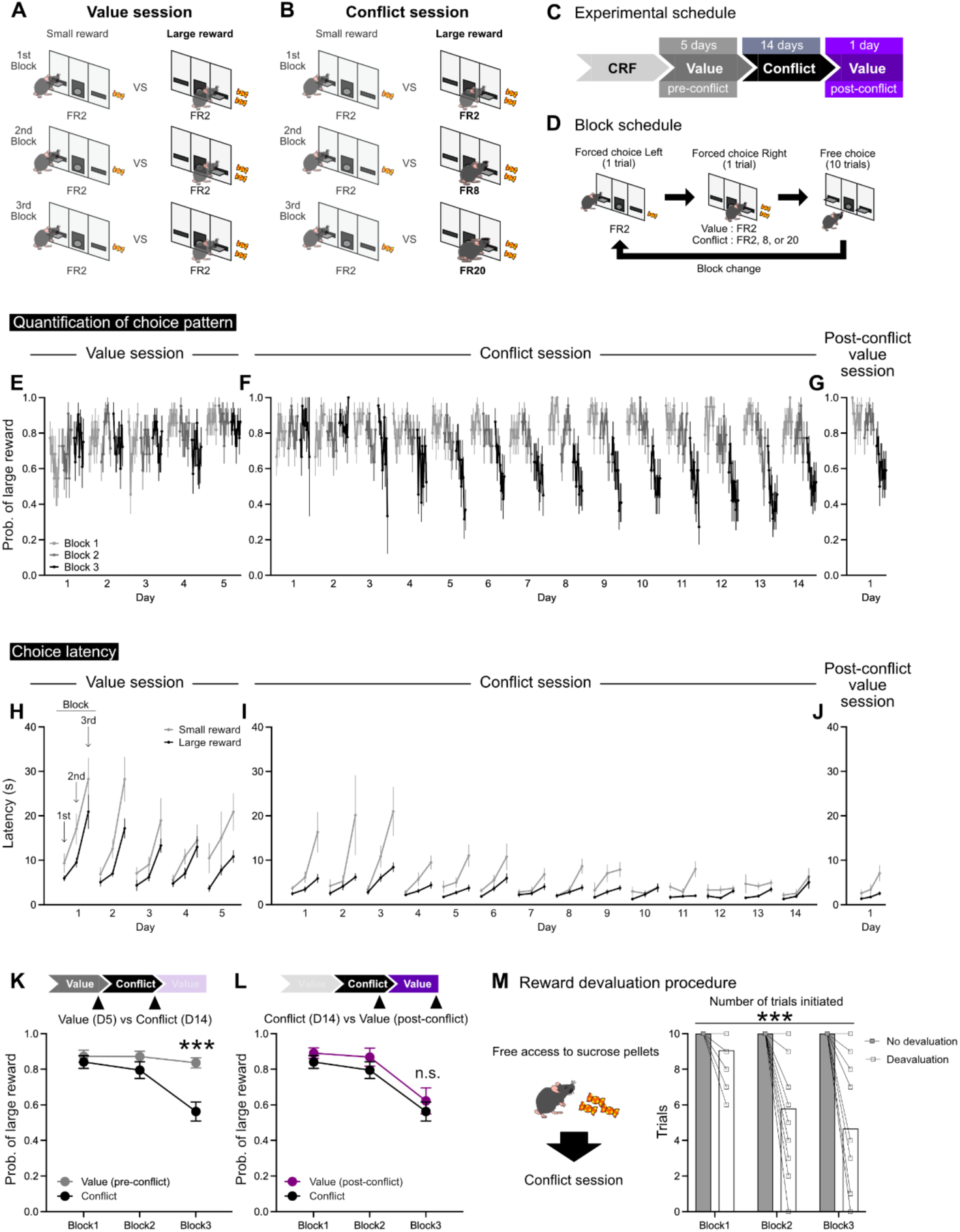
Consolidation of the fixed action schema via a recurrent pattern of goal-directed choice. (A) Diagram of the Value session with equal effort requirements to obtain small and large rewards, asking their capacity to make a reward-value based choice. (B) Diagram of the Conflict session with the varying requirements of efforts to obtain a large reward, asking them to make a conflicting choice between reward sizes and required efforts. (C) Experimental schedule from continuous reinforcement schedule (CRF) to the Post-Conflict Value session. (D) Block transition schedule. (E-G) Probability of large reward choices on each day of value sessions (E), Conflict sessions (F) and one-day Post-Conflict Value session (G) (*N* = 22 mice). (H-J) Choice latency (response latency from lever extension to their choice) when making small or large reward choices on each day of value sessions (H), Conflict sessions (I) and one-day Post-Conflict Value session (J). (K) Probability of large reward choices on Day 5 of the value session (gray) and Day 14 of the Conflict session (black). Two-way repeated measures ANOVA, main effect of Session type: *F*_(1, 42) =_ 9.359, *p* = 0.0039; interaction of Block × Session type: *F*_(2, 84) =_ 8.197, *p* = 0.0006; Sidak’s multiple comparisons: Block 1, *p* = 0.9190; Block 2, *p* = 0.4305; Block 3, *p* < 0.0001. (L) Probability of large reward choices on Day 14 of the Conflict session (black) and Post-Conflict Value session (purple). Two-way repeated measures ANOVA, main effect of Session type: *F*_(1, 42) =_ 1.2, *p* = 0.2766; interaction of Block × Session type: *F*_(2, 84) =_ 0, *p* = 0.9578; Sidak’s multiple comparisons: Block 1, *p* = 0.8620; Block 2, *p* = 0.6692; Block 3, *p* = 0.7845. (M) Reward devaluation procedure and the result of the number of trials mice attempted their choice (*N* = 15 mice). Two-way repeated measures ANOVA, main effect of Treatment: *F*_(1, 28) =_ 24.74, *p* = 0 < 0.0001; interaction of Block × Treatment: *F*_(2, 56) =_ 16.03, *p* < 0.0001; Sidak’s multiple comparisons: Block 1, *p* = 0.6123; Block 2, *p* < 0.0001; Block 3, *p* < 0.0001. Data are presented as mean ± SEM. *p* < 0.05 was considered statistically significant.

After the completion of CRF, we trained animals in Value sessions for five days (Fig. 1A). In the Value session that required an equal effort to obtain large or small rewards, we asked a question of whether mice can make a reasonable choice for a large reward (2 sucrose pellets) than a smaller reward (1 pellet). Indeed, mice appeared to quickly figure out reward sizes associated with each choice on Day 1, and their preference to large rewards became stable across three blocks by the end of Value sessions (Fig. 1E and 1K).

Next, we moved the animals to extensive Conflict sessions in which animals made a conflicting choice between low effort/small reward vs high effort/large reward options under a three-block incremental design of efforts required for large rewards (Fig. 1B, 1D). As expected, they showed a shift in their choice from large to small rewards upon the increment of effort requirements within a daily session (hereafter, the *Large-to-Small* decision shift), and importantly, this shift became a recurrent pattern consistently observed across days, remaining stable in the second half of training sessions (Fig. 1F). A separate analysis of their choice latency indicated the gradual decrease and lesser variability towards the end of Conflict sessions (Fig. 1H-J), suggesting that their choice behavior was temporally stabilized as well. These observations led us to hypothesize that if animals kept choosing in the same way in Conflict sessions, this recurrent pattern of choice, namely the *Large-to-Small* decision shift within a session, would become their fixed rule for action selection. To determine this by an experimental manipulation, we suddenly put the animals back to the original Value session (one-day Post-Conflict Value session) to see whether this recurrent choice pattern persists even after the reward/effort contingency has changed. In fact, mice demonstrated the *Large-to-Small* decision shift much alike to that observed in Conflict sessions (Fig. 1G), even though the efforts for large rewards were no longer increased. Statistical analyses indicate a significant difference between their decisions in the initial Value session and the Conflict session (Fig. 1K), but their choice pattern in one-day Post-Conflict Value session was not significantly different from the Conflict session (Fig. 1L); indeed, it clearly resembles the recurrent choice pattern made in the Conflict session. These findings provide two critical implications. First, it suggests that two-week long Conflict sessions make animals’ choice stereotyped so they acquire a recurrent pattern of choice, that is the *Large-to-Small* decision shift. Second, as this recurrent pattern persists even after having reward/effort contingency changed, animals should have consolidated it into their fixed schema for action selection which then dominates the control of choice, consequently yielding the perseverative responses in the Post-Conflict Value session.

### The recurrent pattern of choice is made under the goal-directed control

One might ask if their recurrent pattern of choice may have become habitual. We thus tested whether their choice behavior is habitual or goal-directed, using a reward devaluation procedure. Using another cohort of mice that had been trained extensively (> 14 days) on Conflict sessions, we gave them free access to sucrose pellets before a Conflict session and tested how many trials they attempted a choice by making the first lever press. We found they significantly reduced the number of attempts in choice (Fig. 1M), suggesting that their behavior is sensitive to reward devaluation so is still goal-directed. Together, we establish an experimental procedure by which mice acquire a recurrent pattern of goal-directed choice and develop it into their fixed schema of action selection by the end of two-week long Conflict sessions.

### Functional screening reveals the orbitofrontal cortex as a candidate brain region responsible for the expression of the recurrent choice pattern

We then sought to identify neural substrates responsible for executing this form of choice. Our observation that mice have consolidated their recurrent choices into the fixed action schema by the end of Conflict sessions (Fig. 1F, 1G and 1L) allowed us to leverage the use of a within-subject design for functional screening. Specifically, we applied chemogenetic inhibition of a brain region or its control on the last two days of Value as well as Conflict sessions in a counter-balanced manner (Fig. 2A), asking a question of how their choice patterns were disrupted. According to literature having suggested possible neural substrates of goal-directed choice behaviors ^11,14,19,40–42^, we selected orbitofrontal (OFC), medial prefrontal (mPFC), anterior cingulate cortex (ACC), nucleus accumbens (NAc), and hippocampus, and injected either AAV-CaMKIIa-hM4Di (OFC, mPFC or ACC) or AAV-hsyn-hM4Di (NAc, hippocampus) into these targeted areas. Among all these inhibitory manipulations in Value and Conflict sessions (Fig. 2B-P), we found that only when OFC activity was inhibited in the Conflict session, animals’ decision pattern was significantly disrupted (Fig. 2D), as evidenced by the absence of the *Large-to-Small* decision shift that had already been formed in Conflict sessions. This finding infers that OFC is a critical locus for this task and allows us to narrow down our focus on identifying the specific role of OFC in the following experiments.

**Figure 2.**
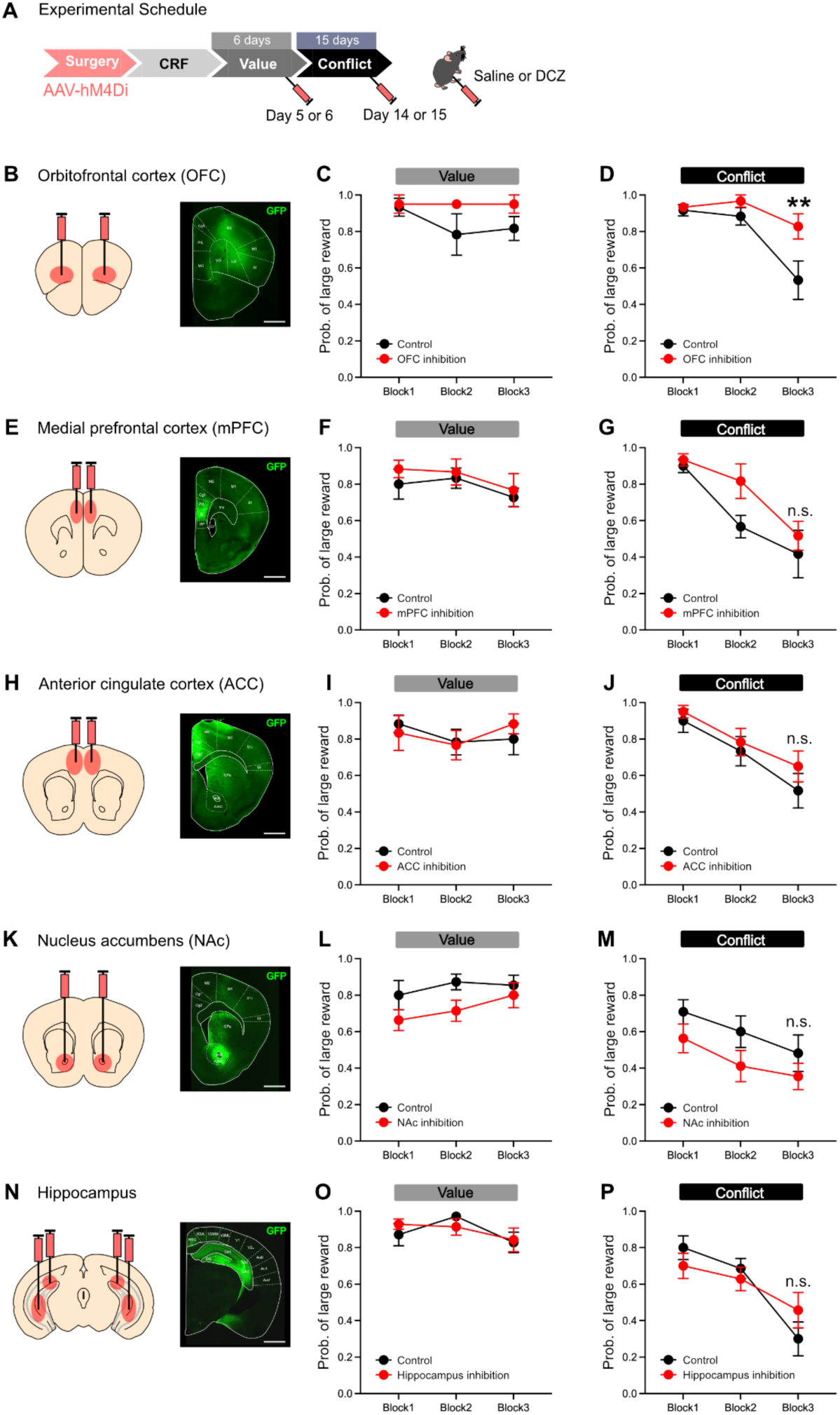
Chemogenetic suppression of OFC disrupts the recurrent choice pattern. (A) Experimental schedule for functional screening by chemogenetic inhibition. (B-D) Surgical strategy to inhibit orbitofrontal cortex and an image of the injection site from an example mouse (B); scale bar, 1 mm. A comparison of results between saline and DCZ administered conditions in value session (C); Two-way repeated measures ANOVA, main effect of Manipulation: *F*_(1, 10) =_ 2.764, *p* = 0.1274; interaction of Block × Manipulation: *F*_(2, 20) =_ 0.949, *p* = 0.4039; Sidak’s multiple comparisons : Block 1, *p* = 0.9971; Block 2, *p* = 0.2183; Block 3, *p* = 0.3985, and in conflict session (D); Two-way repeated measures ANOVA, main effect of Manipulation: *F*_(1, 10) =_ 7.183, *p* = 0.0231; interaction of Block × Manipulation: *F*_(2, 20) =_ 3.129, *p* = 0.0657; Sidak’s multiple comparisons : Block 1, *p* = 0.9961; Block 2, *p* = 0.6902; Block 3, *p* = 0.0039. *N* = 6 mice used. (E-G) Same as above for chemogenetic manipulation of medial prefrontal cortex; scale bar, 1 mm. (F) Results in Value session, Two-way repeated measures ANOVA, main effect of Manipulation: *F*_(1, 10) =_ 0.547, *p* = 0.4765; interaction of Block × Manipulation: *F*_(2, 20) =_ 0.112, *p* = 0.8946; Sidak’s multiple comparisons : Block 1, *p* = 0.7802; Block 2, *p* = 0.9810; Block 3, *p* = 0.9705. (G) Results in Conflict session, Two-way repeated measures ANOVA, main effect of Manipulation: *F*_(1, 10) =_ 2.718, *p* = 0.1302; interaction of Block × Manipulation: *F*_(2, 20) =_ 1.209, *p* = 0.3194; Sidak’s multiple comparisons: Block 1, *p* = 0.9879; Block 2, *p* = 0.1010; Block 3, *p* = 0.7659. *N* = 6 mice used. (H-J) Chemogenetic manipulation of anterior cingulate cortex; scale bar, 1 mm. (I) Results in Value session, Two-way repeated measures ANOVA, main effect of Manipulation: *F*_(1, 10) =_ 0.005, *p* = 0.9468; interaction of Block × Manipulation: *F*_(2, 20) =_ 0.7263, *p* = 0.4960; Sidak’s multiple comparisons: Block 1, *p* = 0.9523; Block 2, *p* = 0.9980; Block 3, *p* = 0.8182. (J) Results in Conflict session, Two-way repeated measures ANOVA, main effect of Manipulation: *F*_(1, 10) =_ 1.086, *p* = 0.3218; interaction of Block × Manipulation: *F*_(2, 20) =_ 0.2784, *p* = 0.7599; Sidak’s multiple comparisons : Block 1, *p* = 0.9528; Block 2, *p* = 0.9528; Block 3, *p* = 0.5174. *N* = 6 mice used. (K-M) Chemogenetic manipulation of nucleus accumbens; scale bar, 1 mm. (L) Results in Value session, Two-way repeated measures ANOVA, main effect of Manipulation: *F*_(1, 20) =_ 3.419, *p* = 0.0793; interaction of Block × Manipulation: *F*_(2, 40) =_ 0.5629, *p* = 0.5740; Sidak’s multiple comparisons : Block 1, *p* = 0.3213; Block 2, *p* = 0.2021; Block 3, *p* = 0.8974. (M) Results in Conflict session, Two-way repeated measures ANOVA, main effect of Manipulation: *F*_(1, 20) =_ 2.694, *p* = 0.1163; interaction of Block × Manipulation: *F*_(2, 40) =_ 0.1387, *p* = 0.8709; Sidak’s multiple comparisons: Block 1, *p* = 0.5188; Block 2, *p* = 0.2960; Block 3, *p* = 0.6246. *N* = 11 mice used. (N-P) Chemogenetic manipulation of hippocampus; scale bar, 1 mm. (O) Results in Value session, Two-way repeated measures ANOVA, main effect of Manipulation: *F*_(1, 12) =_ 0.0099, *p* = 0.9223; interaction of Block × Manipulation: *F*_(2, 24) =_ 0.8882, *p* = 0.4245; Sidak’s multiple comparisons: Block 1, *p* = 0.7990; Block 2, *p* = 0.7990; Block 3, *p* = 0.9957. (P) Results in Conflict session, Two-way repeated measures ANOVA, main effect of Manipulation: *F*_(1, 12) =_ 0, *p* > 0.999; interaction of Block × Manipulation: *F*_(2, 24) =_ 2.319, *p* = 0.1201; Sidak’s multiple comparisons: Block 1, *p* = 0.7327; Block 2, *p* = 0.9342; Block 3, *p* = 0.3864. *N* = 7 mice used. Data are presented as mean ± SEM. *p* < 0.05 was considered statistically significant.

### OFC inhibition does not affect conflicting decisions in the absence of recurrent choice experience

There are two possible interpretations on disrupted decision patterns by OFC inhibition (Fig. 2). First, as we hypothesized, OFC inhibition made animals unable to utilize their learned, fixed action schema, i.e., the *Large-to-Small* decision shift within a session. Alternatively, OFC inhibition simply augmented their choice preference or motivation to large rewards. To untangle these confounding factors, we redesigned a training procedure on Conflict sessions that prevents the fixed action schema from having been formed by precluding recurrent choices to occur (Fig. 3A-C). In the two-week long training period, a session composed of two blocks with varying effort requirements was randomly assigned an ascending or descending order of the effort change between two blocks (Fig. 3A-C). This way disallowed animals to repeat the recurrent use of the *Large-to-Small* decision shift, since here they were supposed to make *Large-to-Small* or *Small-to-Large* decision shifts adaptively, depending on efforts’ increment or reduction in a given session. Animals indeed selected large or small rewards flexibly depending on effort requirements (Fig. 3D). We then chemogenetically inhibited OFC in mice performing the ascending version of the Conflict sessions under a within-subject design, indicating that OFC inhibition did not disrupt their choice pattern (Fig. 3E and 3F). This result suggests that OFC inhibition does not merely alter the patten of conflicting decisions. Furthermore, using another cohort of mice, we measured their motivation to a sucrose reward on the progressive ratio schedule ^43,44^ and found that OFC-inhibited mice showed lower, but not higher, break points (Fig. 3G). This result indicates that OFC inhibition did not augment motivation for rewards. Together, these two observations rule out the possibilities that OFC inhibition enhances their preference to large rewards or increases their motivation; rather, OFC inhibition should have affected the expression of the recurrent pattern of choice, likely by disrupting the capacity to use the fixed action schema for guiding behavior (Fig. 2D).

**Figure 3.**
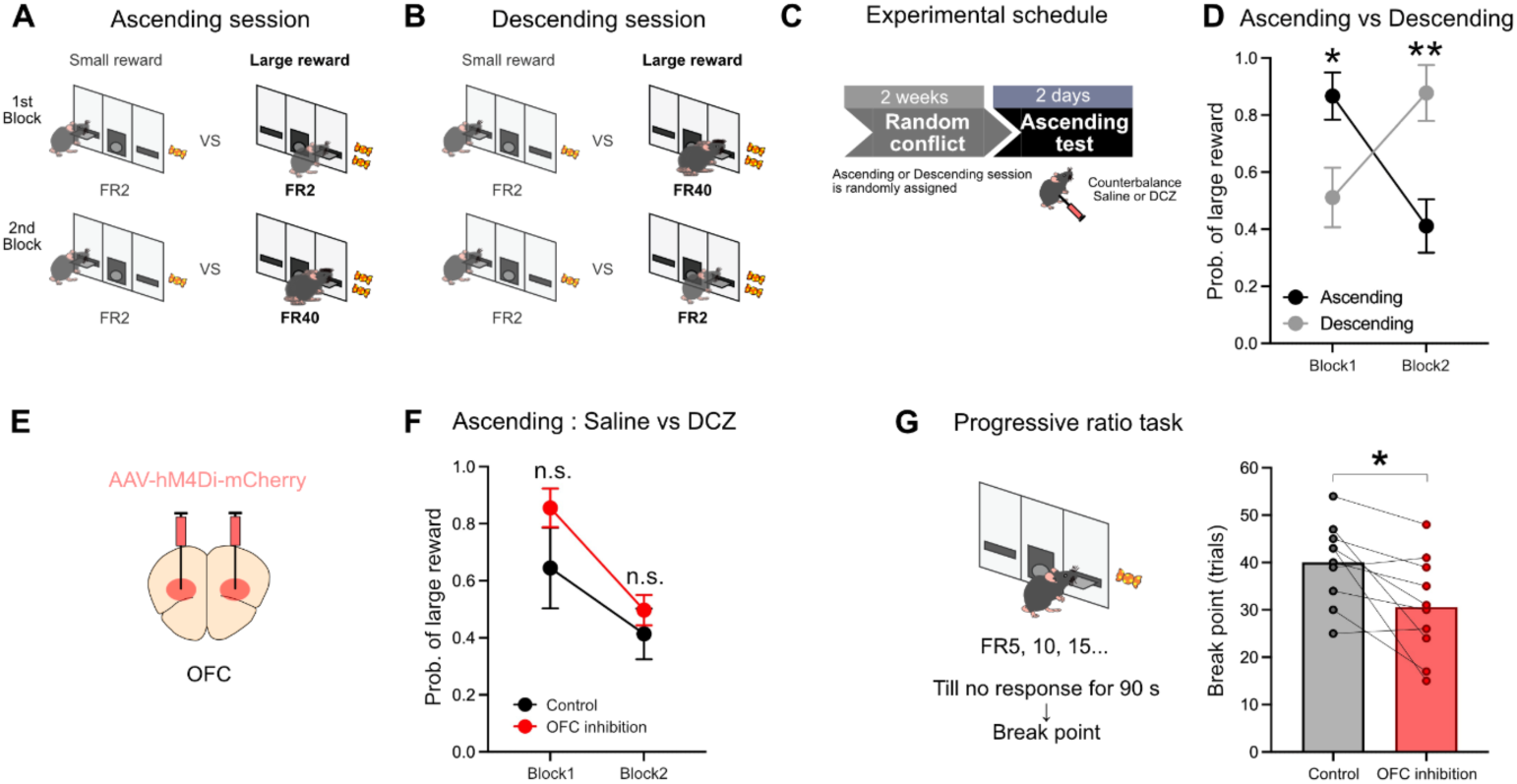
OFC inhibition does not affect conflicting decisions in the absence of recurrent choice pattern. (A-B) Schematic diagrams for order-randomized conflict sessions composed of ascending version (A) that involve the increase in effort requirement and of descending version (B) that is reversed, the decrease in effort requirement. (C) Experimental schedule. (D) A comparison of results between ascending and descending sessions at the end of training (Ascending: black, Descending: gray). Two-way repeated measures ANOVA, main effect of Session type: *F*_(1, 20) =_ 0.3429, *p* = 0.5647; interaction of Block × Session type: *F*_(1, 20) =_ 18.78, *p* = 0.0003; Sidak’s multiple comparisons: Block 1, *p* = 0.0305; Block 2, *p* = 0.0047. *N* = 6 mice used. (E) Surgical strategy to inhibit orbitofrontal cortex. (F) A comparison of results between saline and DCZ administered conditions during the two days of test sessions under the ascending version (Saline: black, DCZ: red). Two-way repeated measures ANOVA, main effect of Manipulation: *F*_(1, 20) =_ 2.454, *p* = 0.1329; interaction of Block × Manipulation: *F*_(1, 20) =_ 0.4601, *p* = 0.5053; Sidak’s multiple comparisons: Block 1, *p* = 0.1281; Block 2, *p* = 0.5371. *N* = 6 mice used. (G) Diagram for the progressive ratio task and the comparison of results between saline and DCZ administered conditions (Saline: black, DCZ: red). Paired *t*-test: *t* _(9) =_ 3.040, *p* = 0.0140. *N* = 10 mice used. Data are presented as mean ± SEM. *p* < 0.05 was considered statistically significant.

### OFC inhibition cancels the perseverative use of the recurrent pattern of choice in Post-Conflict Value session

To directly determine the role for OFC in utilizing the fixed schema for action selection, we designed an experiment under a between-subject design in which OFC was temporarily inhibited in one-day Post-Conflict Value session (Fig. 4A and 4B). A control group that received either AAV-CaMKII-hM4Di with saline to be administered or Sham surgery with DCZ to be administered (Fig. 4C), indicated the stereotypical *Large-to-Small* decision shift shaped by extensive training on the Conflict sessions (Fig. 4D). These mice indeed demonstrated the perseverative use of the fixed action schema in one-day Post-Conflict Value session, as evidenced by nearly identical, recurrent pattern of choice compared to the prior Conflict session (Fig. 4E), which was consistent with the result mentioned earlier (Fig. 1). Similarly, a group of mice to be inhibited OFC also formed the *Large-to-Small* decision shift via Conflict sessions (Fig. 4F and 4G). On the day of Post-Conflict Value session, however, we applied chemogenetic inhibition of OFC and it disrupted their perseverative use of the recurrent choice pattern (Fig. 4H). This result indicates that OFC disruption makes them unable to rely on the fixed action schema and to utilize the recurrent pattern of choice, the *Large-to-Small* decision shift, thus resulting in their choice more adaptive to a change in reward/effort contingency (Fig. 4H). In fact, their choice came back to their original choice pattern alike to the one in the initial Value session (Fig. 4G, gray vs Fig. 4H, red). These results suggest that OFC plays a key role in exploiting the fixed action schema and in exerting the recurrent pattern of choice even upon an abrupt change in contingency, which likely supports behavioral persistency.

**Figure 4.**
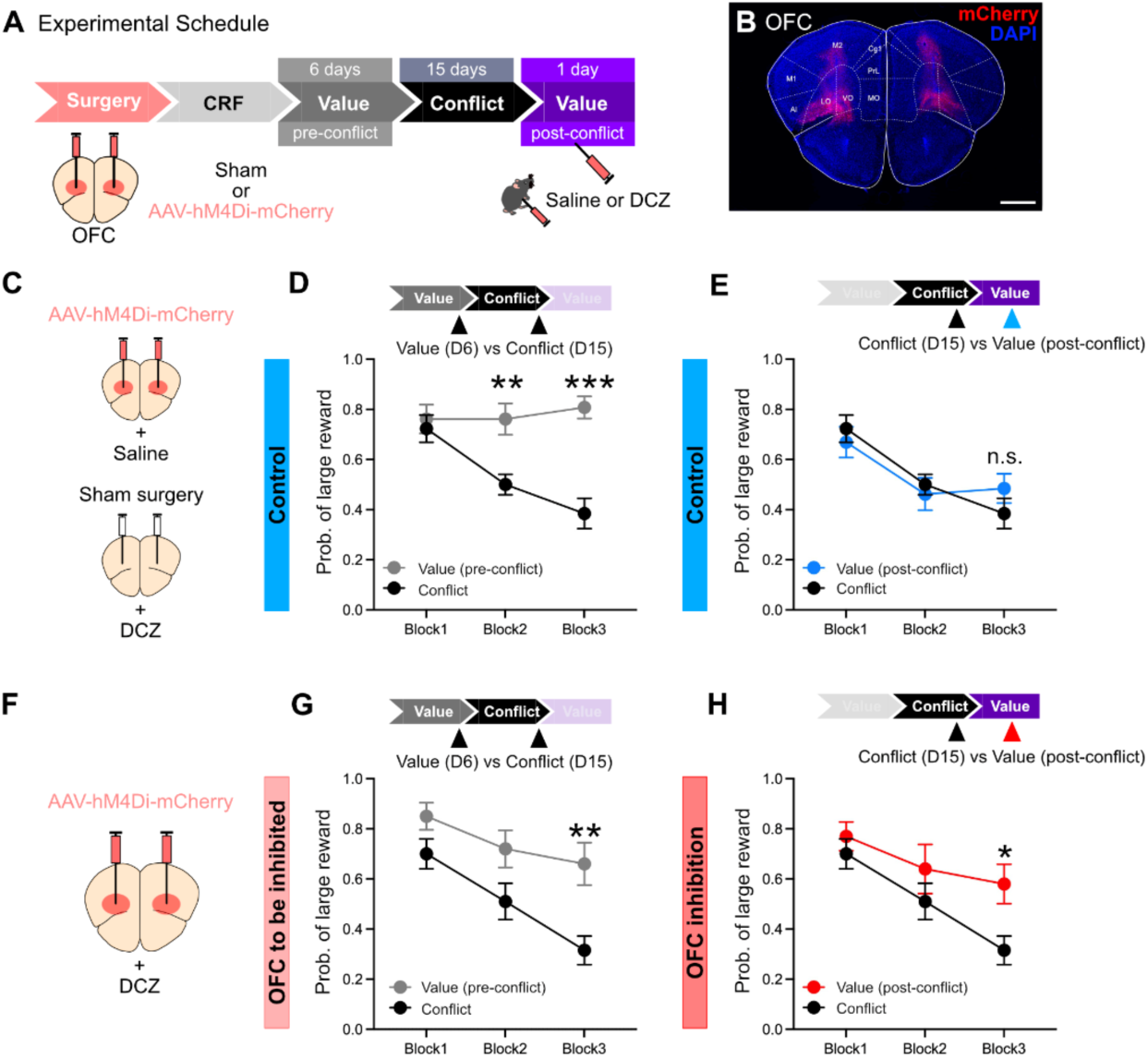
Absence of perseverative choice patterns by transient inhibition of OFC in Post-Conflict Value session. (A) Experimental schedule. (B) A representative image of the injection site in OFC from an example mouse; scale bar, 1 mm. (C) Surgical strategy for the control mice. (D) Probability of large reward choices on Day 6 of Value session (gray) and Day 15 of Conflict session (black). Two-way repeated measures ANOVA, main effect of Session type: *F*_(1, 24) =_ 23.59, *p* < 0.0001; interaction of Block × Session type: *F*_(2, 48) =_ 7.372, *p* = 0.0016; Sidak’s multiple comparisons: Block 1, *p* = 0.9434; Block 2, *p* = 0.0031; Block 3, *p* < 0.0001. (E) Probability of large reward choices on Day 15 of Conflict session (black) and Post-Conflict Value session (blue). Two-way repeated measures ANOVA, main effect of Session type: *F*_(1, 24) =_ 0.0029, *p* = 0.9577; interaction of Block × Session type: *F*_(2, 48) =_ 1.116, *p* = 0.3358; Sidak’s multiple comparisons: Block 1, *p* = 0.8814; Block 2, *p* = 0.9520; Block 3, *p* = 0.5279. *N* = 13 mice used. (F) Surgical strategy to inhibit orbitofrontal cortex. (G) Probability of large reward choices on Day 6 of Value session (gray) and Day 15 of Conflict session (black). Two-way repeated measures ANOVA, main effect of Session type: *F*_(1, 18) =_ 13.65, *p* = 0.0017; interaction of Block × Session type: *F*_(2, 36) =_ 1.278, *p* = 0.2910; Sidak’s multiple comparisons: Block 1, *p* = 0.3276; Block 2, *p* = 0.0958; Block 3, *p* = 0.0021. (H) Probability of large reward choices on Day 15 of Conflict session (black) and Post-Conflict Value session (red). Two-way repeated measures ANOVA, main effect of Session type: *F*_(1, 18) =_ 3.915, *p* = 0.0634; interaction of Block × Session type: *F*_(2, 36) =_ 0.552, *p* = 0.2257; Sidak’s multiple comparisons: Block 1, *p* = 0.8712; Block 2, *p* = 0.5022; Block 3, *p* = 0.0360. *N* = 10 mice used. Data are presented as mean ± SEM. *p* < 0.05 was considered statistically significant.

### OFC ablation leaves conflicting choice intact but disrupts the capability to consolidate the recurrent choice into the fixed action schema

So far, we found that the mice shape their fixed action schema by recurrent execution of similar choices (Fig. 1), which requires intact OFC (Fig. 2). In contrast, OFC inhibition had no effect on conflicting choice itself (Fig. 3), unless the fixed action schema has been formed in the acquisition phase. We also indicated that OFC plays a role in utilizing the recurrent choice even after the contingency change (Fig. 4). These observations led us to two interpretations for the specific role of OFC. First, OFC is critical for expression of the recurrent choice, regardless of the degree to which its action selection is fixated. Second, the OFC consolidates the recurrent pattern of choice into the fixed action schema, which dominates the control of choice in late Conflict sessions and exerts resistance to change in the Post-Conflict Value session. One way to dissociate them is to ablate OFC before behavioral acquisition. If the former is correct, we would not see the recurrent choice pattern in Conflict sessions. If the latter is favored, then OFC ablation would have no impact on the recurrent choice pattern per se, but do have an effect on its consolidation into the fixed action schema, which should lead to the absence of perseveration in Post-Conflict Value session.

We therefore performed excitotoxic lesions of OFC before behavioral experiment and observed the lesion effect (Fig. 5A and 5B). A control group of mice injected with saline indicated their choice preference to large rewards in the initial Value sessions (Fig. 5C), and formed the common *Large-to-Small* decision shift in the Conflict sessions (Fig. 5D and 5I). Importantly, in one-day Post-Conflict Value session, the control group demonstrated the perseverative use of the recurrent choice pattern, as indicated by no statistical difference in the 3^rd^ block of Post-Conflict Value session compared to the Conflict session (Fig. 5E and 5J). This is consistent with the other cohorts (Fig. 1E-G, Fig. 4C-E). OFC-ablated mice were intact in the initial Value sessions (Fig. 5F) and showed the *Large-to-Small* decision shift in Conflict sessions similar to the control (Fig. 5G and Fig. 5K), leaving the acquisition of the recurrent choice pattern intact. Most critically, however, OFC-ablated mice showed rapid updating of their choice pattern in the 3^rd^ block of Post-Conflict Value session that was significantly different from the Conflict session (Fig. 5H and 5L), indicating that their perseverative use of fixed action schema was absent. Indeed, their choice patterns became much more alike those in the initial Value session (Fig. 5K gray line vs 5L red line). Unlike the control or the other cohorts (Fig. 1E-G, Fig. 4C-E) in which the fixed schema of action selection was successfully formed via extensive Conflict sessions, these OFC-ablated mice similarly showed the recurrent pattern of the *Large-to-Small* decision shift but were unable to consolidate it into their fixed action schema. Consequently, their choice was less perseverative upon the contingency change in the one-day Post-Conflict Value session. These findings suggest that OFC plays a specific role in ensuring behavioral persistency by shaping the extensive history of recurrent choices into the fixed action schema to guide behavior.

**Figure 5.**
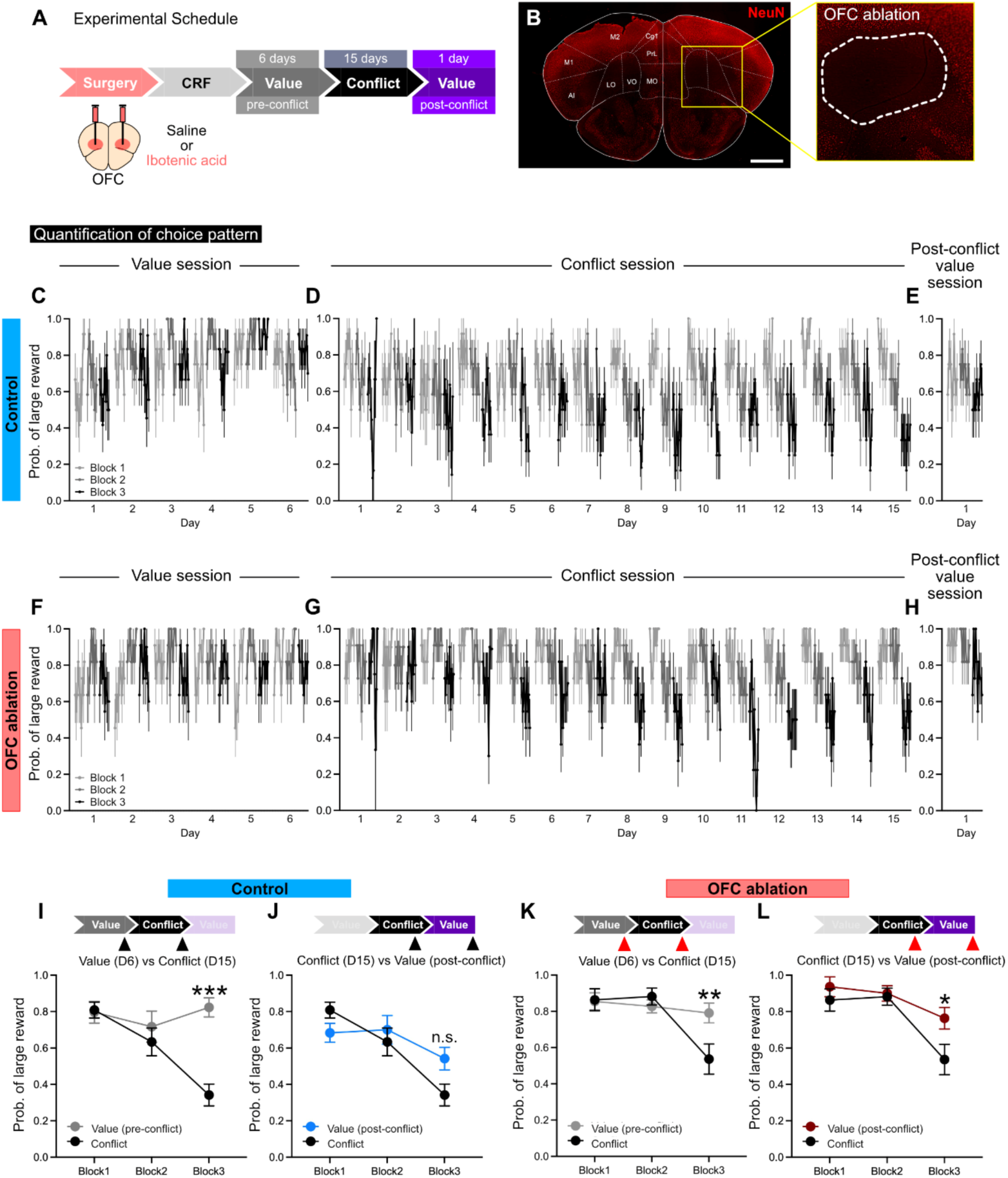
OFC ablation leaves the acquisition of the recurrent choice pattern intact, but disrupts the perseverative choice pattern in the Post-Conflict Value session. (A) Experimental schedule. (B) Representative image of the excitotoxic ablation of OFC in an example mouse; scale bar, 1 mm. An inset indicates the lesioned area labeled by dashed lines. (C-E) Probability of large reward choices in the control group on each day of value sessions (C), Conflict sessions (D) and one-day Post-Conflict Value session (E) (*N* = 12 mice). (F-H) Probability of large reward choices in the OFC-ablated group on each day of value sessions (F), Conflict sessions (G) and one-day Post-Conflict Value session (H) (*N* = 10 mice). (I) Probability of large reward choices in the control group on Day 6 of the value session (gray) and Day 15 of the Conflict session (black). Two-way repeated measures ANOVA, main effect of Session type: *F*_(1, 22) =_ 7.481, *p* = 0.0121; interaction of Block × Session type: *F*_(2, 44) =_ 12.05, *p* < 0.0001; Sidak’s multiple comparisons: Block 1, *p* = 0.9989; Block 2, *p* = 0.7434; Block 3, *p* < 0.0001. (J) Probability of large reward choices in the control group on Day 15 of the Conflict session (black) and Post-Conflict Value session (blue). Two-way repeated measures ANOVA, main effect of Session type: *F*_(1, 22) =_ 0.5236, *p* = 0.4769; interaction of Block × Session type: *F*_(2, 44) =_ 4.747, *p* = 0.0136; Sidak’s multiple comparisons: Block 1, *p* = 0.4222; Block 2, *p* = 0.8416; Block 3, *p* = 0.0839. (K) Probability of large reward choices in OFC-ablated group on Day 6 of the value session (gray) and Day 15 of the Conflict session (black). Two-way repeated measures ANOVA, main effect of Session type: *F*_(1, 20) =_ 1.061, *p* = 0.3153; interaction of Block × Session type: *F*_(2, 40) =_ 7.075, *p* = 0.0023; Sidak’s multiple comparisons: Block 1, *p* = 0.9993; Block 2, *p* = 0.8745; Block 3, *p* = 0.0071. (L) Probability of large reward choices in OFC-ablated group on Day 15 of the Conflict session (black) and Post-Conflict Value session (red). Two-way repeated measures ANOVA, main effect of Session type: *F*_(1, 20) =_ 2.719, *p* = 0.1148; interaction of Block × Session type: *F*_(2, 40) =_ 2.716, *p* = 0.0783; Sidak’s multiple comparisons: Block 1, *p* = 0.7719; Block 2, *p* = 0.9950; Block 3, *p* = 0.0259. Data are presented as mean ± SEM. *p* < 0.05 was considered statistically significant.

## Discussion

Based on an well-established cost-benefit decision task ^38^, we here introduced an experimental procedure in which animals shape the fixed schema of action selection in goal-directed choice. In Conflict sessions, mice executed the recurrent pattern of choice over and over. This extensive, recurrent choice makes their action schema rigidly fixed such that it gradually dominates animals’ choice behavior, consequently yielding the perseverative use of the recurrent choice in Post-Conflict Value session (Fig. 1, Fig. 6A). The following functional screening of brain areas indicated that the OFC inhibition disrupted the choice patterns in the last phase of Conflict sessions (Fig. 2), likely due to their inability to exploit the fixed action pattern (Fig. 6B). In contrast, the same treatment had no effect on animals’ conflicting choice when the fixed action schema was absent (Fig. 3, Fig. 6C). Consistently, the selective OFC inhibition in Post-Conflict Value session made their choice less perseverative upon the change in reward/effort contingency (Fig. 4), which alternatively indicates their inability to exploit the fixated choice pattern (Fig. 6D). Lastly, OFC-ablated mice were intact in executing the recurrent choice pattern itself, but they demonstrated no perseveration to this pattern in the Post-Conflict Value session (Fig. 5), indicating the absence of the capacity in OFC-ablated mice to solidify the recurrent pattern of choice into the fixed action schema (Fig. 6E). Combined all pieces of evidence together, our findings suggest that OFC plays a specific role in ensuring persistency of behavior by consolidating the recurrent pattern of choice into the fixed schema of action selection, which helps guide future choices to use prior experiences via exploitation (Fig. 6F).

**Figure 6.**
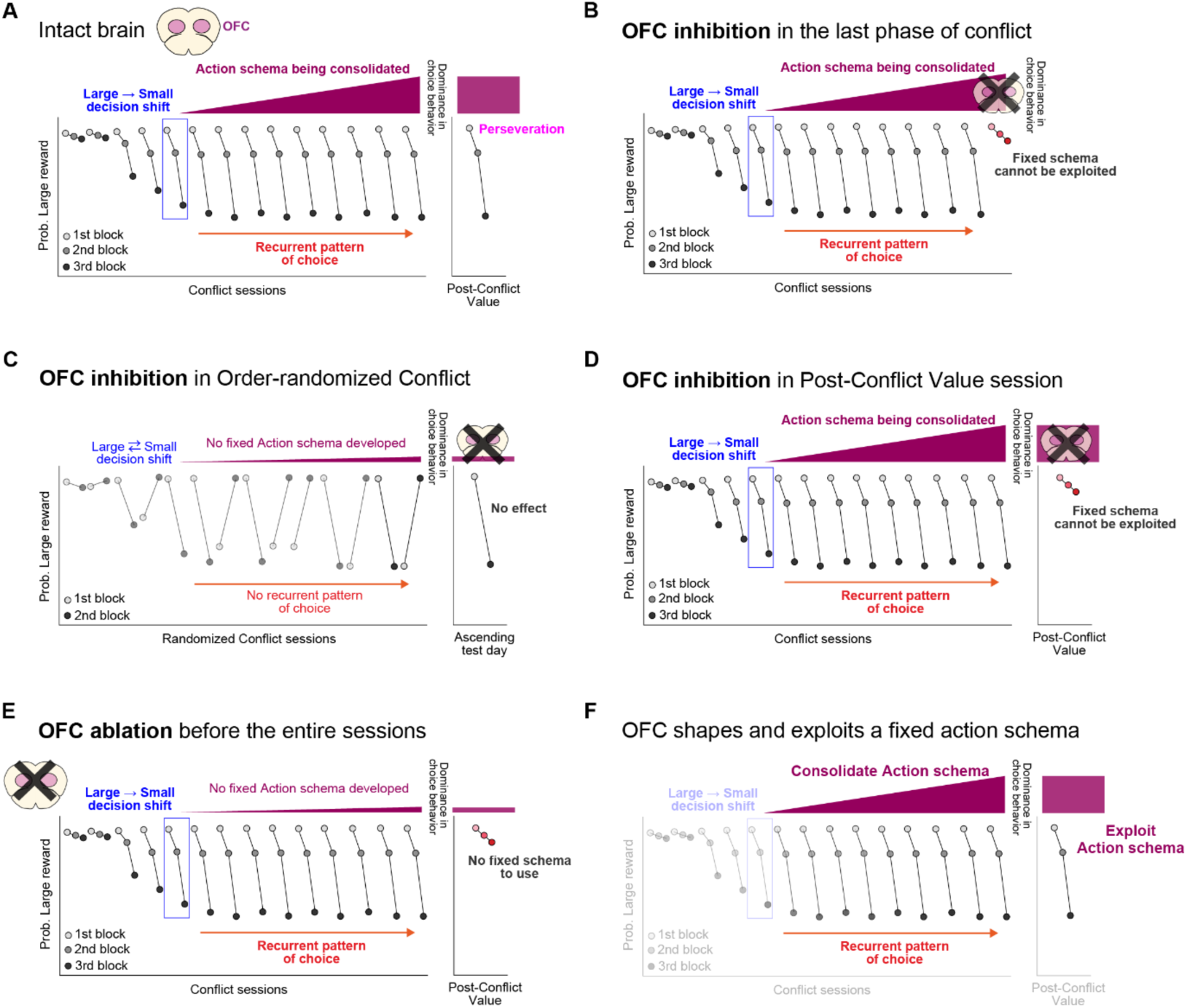
Summary of our findings and interpretations, and proposed roles of OFC. (A) Summary diagram for the formation of the fixed action schema consolidated by the repeated execution of recurrent choices, which then dominates the control of goal-directed choices, thus causing perseveration when the contingency has abruptly changed in the Post-Conflict Value session (Related to Fig. 1). (B) Summary diagram for OFC inhibition in the last phase of Conflict sessions, based on the assumption that OFC plays a role in exploiting the fixed action schema. By the end of Conflict sessions, the fixed action schema dominates the control of choices, so OFC inhibition alters their decision pattern by disrupting the ability to exploit the learned schema of action selection. (C) Summary diagram for OFC inhibition in Order-Randomized Conflict sessions. This paradigm prevents the fixed action schema from being formed, so the OFC inhibition does not affect their choice pattern. (D) Summary diagram for OFC inhibition in Post-Conflict Value session. This session involves an abrupt change in contingency, so mice with intact OFC normally show the perseverative, recurrent choice pattern (Fig. 4), since at this point the fixed action schema dominates their choice. However, without OFC, animals cannot exploit the fixed action schema, resultantly leading to the quick adaptation to the contingency change. (E) Summary diagram for OFC ablation before behavioral acquisition. The absence of OFC leaves intact to make a recurrent pattern of choice characterized by the *Large-to-Small* decision shift, but disrupts the consolidation of this recurrent pattern into the fixed action schema, thus causing the absence of the perseverative choice in the Post-Conflict Value session. (F) Proposed roles for OFC. Given abovementioned observations, it is likely that OFC consolidates a recurrent pattern of choice into the fixed schema of action selection and exploits it when making routine-like choice.

The framework that distinguishes habitual vs goal-directed control of actions has provided a foundation to the understanding of animals behaviors ^8,12,13^. Yet, some others have argued a more intricate way of behavioral control that explains routines, sequencing or schema of actions, positing that animal behaviors can be stabilized, chunked and sequenced efficiently in a goal- oriented manner ^11,28,29,34^, which are dissociable from the definition of habits. Our study provides a behavioral procedure in the context of decision that allows us to test animals’ ability to shape their repeated choice patterns into the fixed action schema and investigate its neural mechanisms. Importantly, previous studies on which the concept of stereotypical but goal-directed actions was founded have mainly focused on action sequencing and chunking of multiple motor responses ^21,34,35^. Given this background, our findings have added another layer of behavioral persistency at a decision level. Indeed, psychological studies have proposed that humans often make perseverative decisions in experimental ^45–48^ and more natural settings ^49,50^. Taking these into account, we here offer an experimental approach using a mouse model to interrogate neural mechanisms for consolidation of the fixed action scheme for decisions.

Our results indicate OFC as a critical neural locus that shapes the recurrent choices into their fixed action schema for decision making. Many pieces of evidence in animal and human studies suggest that OFC is involved in flexible decision making ^7,51–57^, representations of action values ^51,58^, and model-based reinforcement learning ^7,59,60^. In addition, OFC neurons are found to be more correlated with goal-directed responding than habitual in instrumental paradigms of mice, suggesting that OFC is critical for shifting action control towards a goal-directed manner ^61^. While this is consistent with our result that OFC is necessary for goal-directed decision, it contradicts with our core finding that OFC solidifies the repeated pattern of choice into the fixed action schema that enables the persistency in behavioral outputs. Notably, Groman et al. have suggested that OFC plays multi-directional roles in decision making processes in rats, indicating that OFC → nucleus accumbens pathway makes their choice flexible while OFC → amygdala stabilizes the learned value of action so yields behavioral persistency ^7^. In a similar vein, OFC → secondary motor cortex projections have been identified to be necessary for exploitation of learned actions ^62^. Taking these findings together with ours, we assume that in a specific context of the decision paradigm involving extensive history of recurrent choices, OFC is capable of exploiting learned values of actions based on prior experiences and consolidates such recurrently performed choice into a fixed action schema to guide goal-directed behavior. To conquer the limitation of our study such as generalizability of OFC-derived action schema in various contexts and the lack of circuit and synaptic mechanisms, we urge following studies that investigate neural representations of the fixed action schema by OFC neurons, identify critical upstream and downstream of the OFC-centered circuitry, and test how much generalizable the role of OFC in shaping the action schema is by the use of repetitive exposure to many other decision scenarios.

Lastly, our findings provide a valuable observation in psychiatry such as addiction and obsessive-compulsive disorder that involve exaggerated repetition of stereotyped behaviors and maladaptive decisions ^20,63–65^. Given the role for OFC in shaping recurrent choices into their fixed action schema, this mechanism may adversely promote the development of their behavioral addictions for drug seeking, substance abuse and compulsion, all involving the lack of behavioral inhibition. Elucidating whether and how OFC’s mechanisms for consolidating the fixed action patterns play a role in the acquisition process of such maladaptive behaviors will be of value to translational research. In summary, we here propose that OFC plays a crucial role in consolidating an action schema by recurrent goal-directed choices, giving insights into how animals acquire persistency and stability of action and how this ability can be compromised with diseases.

## Acknowledgement

We would like to thank Drs. Jane R. Taylor, Carlos Alos-Ferrer, Susumu Setogawa, Keiji Ota for scientific discussion and fruitful suggestions. We also thank all the members of Abe Laboratory at Tohoku University for their help and discussion on the present study.

## Author Contributions

Conceptualization: R.T, and S.A; Animal surgery: R.T; Behavioral experiments and analysis: R.T; Histology: R.T; Resources: S.A and K.A; Funding: S.A and K.A; Supervision: S.A and K.A; Writing an original draft: S.A; Editing and Finalizing the manuscript: R.T, S.A, and K.A.

## Data availability

Data and materials are provided upon a reasonable request to the corresponding author.

## Materials and Methods

### Animals

We used adult male mice (C57BL/6J) obtained from SLC Japan and those bred in our animal facility. Mice were housed in plastic cages in a temperature- and humidity-controlled room under a 12-h light/dark cycle (light on at 8:00 a.m. and off at 8:00 p.m.) and could access standard food and water *ad libitum*. The care and experimental procedure of animals were reviewed and approved by the Institutional Animal Care and Use Committee of Tohoku University. All experiments and maintenance were performed following relevant guidelines and regulations.

### Behavioral procedure

All behavioral experiments are performed in an operant conditioning chamber that contains two receptable levers, house light, sucrose pellet dispenser, and pellet magazine (Med Associates Inc.). A sucrose pellet was used as a food reward (20 mg, Dustless Precision Pellets^®^, Bio-Serv). Behavioral events were recorded and saved by the Med Associates’ software (MED-PC IV). During behavioral experiments, mice were food-restricted at approximately 85% of their body weight.

#### Continuous reinforcement schedule (CRF)

CRF was performed under a fixed-ratio schedule of reinforcement (FR1) in which each lever press delivered a sucrose pellet. Mice were given a maximum of 15 rewards on each left or right lever for initial three days, and were given a maximum of 30 rewards on each lever for another three days. The order of left or right lever presentation was pseudorandomized. A daily CRF session ended by either the maximum number of rewards obtained or 30 minutes passed.

#### Reward-Value based choice session (Value session)

A daily session consisted of three blocks. Each block began by a forced choice trial on each of small and large reward options to instruct a mouse on the size of a reward and effort. First, only the left lever associated with a small reward was presented to the mouse and when completed two lever presses (FR2), a sucrose pellet was given. Next, only the right lever associated with a large reward was presented and upon the completion of FR2, two sucrose pellets were given. In the following 10 trials of free choice, mice are presented both left and right lever to make a choice and after they chose one of the levers, the other was retracted. After having 10 trials of free choice completed or 10 minutes passed in each block, the session moved onto the next block. In this Value session, there was no change in effort requirement across session. Inter-trial interval was set at 20 seconds. If mice did not complete required lever presses for 25 seconds after their choice, the lever they chose was also retracted, no reward was given and the trials were scored as an omission. Mice were trained on value sessions for 5–6 days before the commencement of the following Conflict sessions. Post-Conflict Value session was performed the same way as above after extensive training of Conflict sessions.

#### Reward-Value/Effort conflict session (Conflict session)

Similar to above, a daily session consisted of three blocks. Each block began by a forced choice trial on each of small and large reward options to instruct a mouse on the size of a reward, followed by 10 trials of free choice. In this Conflict session, the required number of lever presses for small rewards remained unchanged at FR2 but it increased for large rewards in proportion to the progression of blocks (Block 1: FR2, Block 2: FR8, Block 3: FR20). At the initial phase of training, there was no constraint of how long they took to complete the required lever presses without omission trials for three days (all the groups with chemogenetic manipulations or neural ablation) or seven days (wild-type mice in Fig. 1). After this phase, we then set 25 seconds for mice to complete the required number of lever presses; otherwise the lever they chose was retracted, no reward was given and the trial was scored omission. Mice were trained on the Conflict sessions for 14–15 days before moving onto the next Post-Conflict Value session.

#### Order-randomized Conflict sessions

This session consisted of two blocks. One block required equal amounts of efforts (FR2) to obtain a small or large rewards, while the other block asked conflicting decisions with FR2 for small and FR40 for large rewards. At each block, we prepared twice of forced choice trials on each of small and large reward choices, followed by 15 trials of free choice. Importantly, the ascending version of the session asked animals to choose small or large rewards with equal efforts in the 1^st^ block and to make conflicting decisions in the 2^nd^ block. The descending version was its reversal such that mice performed conflicting decisions in the 1^st^ block, followed by the 2^nd^ block where they chose small or large rewards with equal efforts (Fig. 3). We randomly assigned either the ascending or descending session on a daily basis, thus asking mice to make an adaptive choice in each given session, which prevents them from fixing their choice patterns, i.e., the *Large-to-Small* decision shift as seen in the other behavioral schedule (Fig. 1). On two consecutive days of chemogenetic manipulations, we only assigned the ascending version to all the mice. Inter-trial interval was set at 20 seconds, and when not completing the lever presses within 25 seconds, a reward was omitted and the trial was scored omission.

#### Behavioral analyses in Value, Conflict and Order-randomized Conflict sessions

Behavioral data were analyzed offline using custom MATLAB codes. Probability of large reward choices was calculated by the number of large reward choices divided by the total number of trials animals performed in each block. This probability of large reward choices was also calculated by a single-trial basis (Fig. 1E-G and Fig. 5C-H), where the single plot indicates the mean of large reward choice of all mice at each trial. Choice latency from lever presentation to their choice was also measured in each block.

#### Progressive ratio task

Using only a lever presented to animals, we asked them to complete the required number of lever presses for a sucrose pellet. The required number was proportionally increased by five times starting from 5 presses until the time at which no lever press had occurred for the last 90 seconds or 60 minutes passed in the session. The last trial they completed the required lever press was recorded as a break point for quantification of their motivation to a sucrose reward. Inter trial interval was set at 20 seconds.

#### Reward devaluation procedure

To determine whether animals’ choice is sensitive to reward value, reward devaluation was applied. Mice were given free access to sucrose pellets in their home cage for an hour before the Conflict session. In this procedure, we compared the number of trials they initiated in each block between non-devalued condition (prior session to devaluation procedure applied) and devalued condition.

### Surgical procedure

Mice were placed on a stereotaxic frame and anesthetized by inhalation of isoflurane (3–4% induction and 1–2% maintenance) in a sterilized condition. For comprehensive screening (Fig. 2), we used either a 1:1 mixture of AAV2/9-Syn-EGFP and AAV2/9-hSyn-hM4Di-flag or AAV2/9-CaMKIIa-hM4Di-2a-EGFP-WPRE (these viruses made in house as reported previously ^66^) for chemogenetic inhibition experiments and applied pressure injections by a 33-gauge needle attached to a Hamilton syringe. We bilaterally injected the mixture into nucleus accumbens (0.3 µl injected at each site, AP: +1.3, ML: 1.2, DV: 4.2/3.8, all from bregma or dura), or dorsal and ventral hippocampus (0.3 µl injected at each site, AP: –2.0, ML: 1.8/0.8, DV: 1.6 and AP: –3.3, ML: 2.9, DV: 3.3/2.0). We also bilaterally injected the AAV2/9-CaMKIIa-hM4Di-2a-EGFP-WPRE in orbitofrontal cortex (0.5 µl injected at each site, AP: +2.3, ML: 1.3, DV: 1.8), into medial prefrontal cortex (0.15 µl injected at each site, AP: +1.7, ML: 0.4, DV: 1.8/1.4/1.0), or into anterior cingulate cortex (0.3 µl injected at each site, AP: +1.5, ML: 0.5, DV: 1.1 and AP: +0.8, ML: 0.5, DV: 1.1). In the other experiments for chemogenetic inhibition of OFC, we injected AAV9-CaMKIIa-hM4Di-mCherry (Addgene viral prep #50477-AAV9, a gift from Bryan Roth) into bilateral OFC (0.5 µl injected at each site, AP: +2.3, ML: 1.3, DV: 1.8). We also performed sham surgery in which the same surgical procedure without virus injections was conducted. For neuronal ablation by excitotoxic lesions, we bilaterally injected 0.4 µl of ibotenic acid (10 mg/ml in saline, Sigma-Aldrich) into OFC (AP: +2.3, ML: 1.3, DV: 1.8). For a control group, we bilaterally injected 0.4 µl of saline into OFC. Upon each injection, the needle was left in place for another 5 min to allow for the solution to spread. After surgery, animals were monitored for signs of stress or discomfort. The mice that underwent surgery had at least a week of recovery and subsequently proceeded to behavioral experiments.

### Chemogenetic inhibition in behavioral experiments

For chemogenetic inhibition experiments under the within-subject design, we applied chmogenetic inhibition by intraperitoneal administration of deschloroclozapine (DCZ, 0.1 mg/kg dissolved in DMSO) or saline mixed with DMSO, 30 minutes prior to behavioral testing. This comparison was made in a counterbalance manner for two testing days. For those under a between-subject design, we similarly administered DCZ or saline to different cohorts of mice 30 minutes prior to the behavioral testing.

### Histology and microscopy

After the completion of behavioral experiments, animals were euthanized and perfused by 4% paraformaldehyde (PFA, Nakarai-tesque), and extracted brains were kept in 4% PFA in the refrigerator (4°C). These brains were then kept in 30% sucrose for > 24 hours before cutting and sectioned coronally at 50 µm using a freezing microtome (REM-710, Yamato kohki Industrial, Co., LTD.). Serial sections were collected and divided into four numbered vials. Selected vials were processed with an interval of 200 µm (1 out of 4) for immunohistochemistry. Sections were first rinsed with phosphate-buffered saline containing 0.9% NaCl (PBS) for 30 minutes, and put in PBS+, i.e. PBS containing 2% normal horse serum and 0.5% Triton X-100 for 1 hour. For visualization of eGFP signals, the sections were then incubated overnight at room temperature in an anti-GFP polyclonal antibody raised in rabbit (Cat#598, MBL Life Science) diluted at 1:1000 in PBS+. Subsequently, the sections were incubated in secondary Anti-rabbit IgG-Alexa Fluor 488 (Cat#A-32731, Thermo Fisher Scientific) at 1:450 for 2 hours. Similarly, we also performed immunohistochemical staining against NeuN by an anti-Fox3 antibody raised in mouse (Cat#SIG-39860, Biolegend, 1:1000) followed by anti-mouse IgG-Alexa Fluor 555 (Cat#A-21422, Thermo Fisher Scientific, 1:450). Upon the completion of all the steps above, the sections were cover-slipped by mounting medium (FluorSave Reagent, Cat#345789, Merck Millipore) with DAPI (D523, Dojindo, 1:1000) for microscopy. Representative fluorescent microphotographs were obtained by a digital camera attached to a Keyence microscope (BZ-9000), followed by visualization of images processed by Image J software.

### Statistical analyses

Statistical analyses were performed using a software GraphPad Prism 7. For the analysis of probability of large reward choice and number of trials in reward devaluation procedure, we performed Two-way repeated measures ANOVA followed by Sidak’s multiple comparisons. For the analysis of progressive ratio schedule, we performed paired t-test. *p* < 0.05 was considered significant.

